# Asymmetrical myofiber architecture along the murine tibialis anterior suggests distinct functional regions

**DOI:** 10.1101/2020.01.17.910422

**Authors:** Davide Bindellini, Lenard M. Voortman, Cyriel S. Olie, Maaike van Putten, Erik van den Akker, Vered Raz

**Author notes:** Postal address: Albinusdreef 2, 2333 ZA Leiden, The Netherlands. **Key points summary** - The study demonstrates that a bipennate muscle, such as tibialis anterior, has an asymmetric architecture along the entire muscle. - Based on geometrical features of myofibers, two main muscle regions could be defined. - The proportions of muscle physiological features, such as myofiber-type and neuromuscular distribution, highly correspond to the defined muscle regions. - Based on our findings, we recommend that the entire muscle should be considered in future studies that assess muscle physiology in disease and aging conditions.

## Abstract

Skeletal muscle function is inferred from the spatial arrangement of myofiber architecture and the molecular and metabolic features of myofibers. Features of myofiber types can be distinguished by the expression of myosin heavy chain (MyHC) isoforms, indicating contraction properties. In most studies, a local sampling, typically obtained from the median part of the muscle, is used to represent the whole muscle. It remains largely unknown to what extent this local sampling represents the entire muscle. Here we studied myofiber architecture over the entire wild type mouse tibialis anterior muscle, using a high-throughput procedure combining automatic imaging and image processing analyses. We reconstructed myofiber architecture from consecutive cross-sections stained for laminin and MyHC isoforms. The data showed a marked variation in myofiber geometric features, as well as MyHC expression and the distribution of neuromuscular junctions, and suggest that muscle regions with distinct properties can be defined along the entire muscle. We show that in these muscle regions myofiber geometric properties align with biological function and propose that future studies on muscle alterations in pathological or physiological conditions should consider the entire muscle.

## Introduction

Skeletal muscles facilitate mobilization and stability of the skeleton, which is highly determined by muscle architecture. Muscle architecture is broadly defined as the organization of muscle fibers (myofibers) relative to the axis of force generation. Muscle architecture is broadly classified into three types: 1)- myofibers arrangement parallel to the force-generating axis on both side of the muscle is called bipennate; 2)- myofibers that are oriented at a single angle relative to the force generating axis is termed unipennate; and 3)- multipennate architecture describes most muscles where myofibers are oriented at several angles relative to the axis of force generation (Otten, 1988; Lieber & Fridén, 2000; Lieber Richard & Ward Samuel, 2011). The architecture of parallel myofibers is mostly suitable to produce extensive force and stabilization. In addition to myofiber arrangement, contraction properties are also determined by myofiber typing. The fast- or slow-twitch myofibers are the two main types, which can be distinguished by metabolic and molecular networks (Schiaffino & Reggiani, 2011; Baskin *et al.*, 2015; Lang *et al.*, 2018; Raz *et al.*, 2018b). Myofibers contractile properties are marked by the expression of myosin heavy chain (MyHC) isoforms (Schiaffino & Reggiani, 2011). The fast-twitch myofibers express MyHC-2A, -2B and -2X isoforms, slow-twitch express MyHC-1. The MyHC-2B type myofibers produce energy by anaerobic glycosylation, whereas the MyHC-2A type myofibers are oxidative and use fatty acids for energy production (Schiaffino & Reggiani, 2011). Alterations in MyHC isoform expression occur in physiological conditions like training or aging, and in pathological conditions (Pette & Staron, 2000). Staining for MyHC isoforms combined with high-throughput image quantification advanced the understanding of myofiber composition and myofiber transition in disease and age(ing) conditions (Riaz *et al.*, 2015; Riaz *et al.*, 2016; Raz *et al.*, 2018a). Studies on myofiber composition and changes in disease animal models are carried out in cross-sections taken from a median region of the muscle. How the entire muscle changes in disease and aging has not been studied.

So far, modelling of myofiber architecture was mainly studied with non-invasive imaging procedures which do not discriminate between myofiber types (Damon *et al.*, 2011; Sullivan *et al.*, 2019). Therefore, the impact of myofiber typing on myofiber architecture and muscle function is not fully elucidated. Here we report a proof-of-principle study in the tibialis anterior (TA) muscle, where we investigated myofiber architecture and myofiber-typing over the entire muscle. The TA muscle connects the knee to the foot, mobilizing the lower leg in a single axis (Mathewson *et al.*, 2012). The mouse TA has a longer sagittal (longitudinal) axis over the frontal and vertical axes. Myofiber contraction in the TA is stimulated by an action potential generated in the neuromuscular junction (NMJ). The distribution of NMJs along the muscle would inferred on contraction potential (Lomo, 2003). The spatial distribution of NMJs along the muscle is not sufficiently studied.

Here We modelled myofiber architecture using myofiber-typing over the entire muscle, and compared our results to imaging-based analysis of TA architecture (Heemskerk *et al.*, 2005; Lovering *et al.*, 2013; Moo *et al.*, 2016). We show that myofiber architecture is not uniform along the muscle. Combining myofiber morphological features with myofiber-typing and NMJs distribution we suggest that a TA muscle is composed of three regions, which together form the muscle spatial layout and functionality.

## Materials and methods

### Mouse and immunohistochemistry procedures

The TA muscles were collected from two 8-weeks-old male C57BL/6J wild type mice. Each muscle was sectioned in a transverse or longitudinal axis. In total, 603 transverse sections of 10 μm thick, covering the entire muscle were generated with the CM3050-S cryostat (Leica Germany)(Moo *et al.*, 2016). Sections were pasted on “PTFE” printed slides, 12 well – 5mm diameter, (Electron Microscopy Sciences, USA). Sectioning of the entire muscle was performed in a single day, avoiding any differences in the cutting angle and batch effect. From the TA muscle of the second mouse, longitudinal sections were generated. Muscle sections were stored at −20°C prior to staining.

The immunohistochemical procedure was performed with a staining protocol that is detailed in (Raz *et al.*, 2018a). Primary antibodies that were used in this study are: Rabbit-anti-Laminin (1:1000, Abcam, UK). Secondary antibodies used are: Goat-anti-Rabbit_Alexa-647 (1:1000, Life Technologies, USA), Goat-anti-Mouse_Alexa-488 (1:1000, Life Technologies, USA). Fluorescently conjugated compounds: antibodies to MyHC-2B_ Alexa fluor-488-conjugated (1:400; [11), MyHC-2A_ Alexa fluor-594-conjugated (1:1000; [11), α-Bungarotoxin_Alexa fluor-488 conjugated (1:1000; Thermo Fisher). Slides were mounted with ProLong Gold (Invitrogen, USA). All cross-sections were simultaneously stained using a single antibody mix, in order to eliminate variations between staining session.

The length of the entire muscle or individual myofibers was measured from longitudinal sections.

### Imaging

Imaging was performed with the Pannoramic 250 Flash III slide scanner (3DHISTECH, Hungary). Imaging of all cross-sections was carried out in a single session, eliminating batch effects. Imaging of the longitudinal sections was carried out using the same Pannoramic 250 Flash III slide scanner. Imaging of α-Bungarotoxin signal was also performed using a Leica DM5500. Manual counting of α-Bungarotoxin signal in NMJs was carried out using a digital magnification where the NMJ structure could be easily recognized (Fig. S4).

### Image processing, quantification and alignment

Slide scanner image files were converted to TIFF using CaseViewer (3DHISTECH, Hungary). After conversion, sections were sorted, selected, cropped and scaled using custom MatLab (MathWorks Inc.) scripts. The selection step, combined with allowing gaps of 150 μm or less, reduced the number of sections from 605 to 150. Scaling of the images (down-sampling) is essential to decrease images size and resolution and thereby increase the processing speed. All sections were exported at a lower resolution scale.

Myofiber segmentation was carried out with the laminin staining using the MuscleJ macro in ImageJ (Mayeuf-Louchart *et al.*, 2018), after following modifications: the image pre-processing step (prior to segmentation) was changed to match our image resolution, implementation of an automatic loading of the whole image set, and an automatic saving of the segmented mask. Poorly segmented images were removed from further analysis. Every segmentation was visually compared to the stained image and subsequently 21 sections were excluded from further analyses due to improper segmentation. This exclusion did not affect the maximum distance allowed between two sections (150 μm). The remaining 129 sections were then mapped to their physical position along the muscle longitudinal axis. Section position was calculated with the first section at 0 µm and consecutively increasing by 10 µm (sections thickness), including the position of the excluded sections. Median and variance for both CSA and circularity were calculated per section. MyHC MFIs were obtained after background correction that was separately applied to each image and each fluorophore. Additionally, a ‘2A mask’ was generated marking the 2A-positive fibers, which were added onto the laminin-segmentation mask. Fibers were considered 2A-positive when their MFI was greater than the MFI mean + one standard deviation of all the fibers per section. The alignment of the images was performed with the ‘2A masks’ in ImageJ using Linear Stack Alignment with SIFT (scale-invariant feature transformation) (Lowe, 2004). This method aligns sections consecutively one by one.

Myofiber composition analysis was carried out using R (version 3.5.1). We first excluded fibers that were not properly segmented. In total 153930 fibers were obtained from the eligible 129 cross-sections. MFI values were scaled per section in order to reduce technical differences between the sections, and analysis was carried out after a natural logarithm (ln) transformation of MFI values. Smoothing lines were calculated and plotted using the smooth.spline function in R, using smoothing parameter=1. Statistical significance was assessed with the Pearson’s chi-squared test.

## Results

**To** develop a proof-of-principle protocol for reconstruction of the 3D-architecture of an entire muscle, we selected the tibialis anterior (TA) of a wild type mouse for its small size (dimension 6×2×2 mm; Fig. 1A), which is suitable for such an initial study. To avoid batch effects, each step in the protocol: sectioning, staining or imaging was completed within a single day per each step. Together, this procedure minimizes technical variation that could give rise to potential confounding batch effects. Imaging and image processing were carried out using high-throughput automatic procedures. In this study two muscles from different animals, same age and gender, were used. One muscle was cross sectioned from the proximal to the distal end, and the other was sectioned on the longitudinal axis. Image processing and quantification were carried out on the cross-sections, and the longitudinal sections used as reference for coordinates and validations (Fig. 1B). The axes of the TA muscle were defined as, proximal and distal on the longitudinal axis connecting knee to foot, respectively, and on the transverse axis posterior or interior marks the position relative to the tibia (Fig. 1C). Myofiber contours were stained with an anti-laminin antibody, and myofiber types were recognized with an antibody mix to MyHC-2A and MyHC-2B (Fig. 1C and 1D). From the MyHC isoforms that are commonly used for myofiber typing in mouse (Bloemberg & Quadrilatero, 2012), MyHC-2B is most prominently expressed in TA muscle and MyHC-2A is expressed in only a subset of myofibers (Riaz *et al.*, 2015). Throughout the entire muscle, myofibers positive to MyHC-2B were more prominent compared with MyHC-2A (Fig. 1D). Myofibers expressing MyHC-2A were measured to be about 160-250 μM in length whereas, MyHC-2B myofibers encompassed the entire muscle length (about 6 mm) (Fig. 1C). Immunofluorescence of the longitudinal and cross-sections confirmed asymmetry in myofiber type distribution: MyHC-2A myofibers were mainly found at the interior side, and major architectural differences between distal and proximal sides (Raz *et al.*, 2018a). Moreover, we observed differences in myofiber orientation over the longitudinal axis: at the proximal side, myofiber length was parallel to the longitudinal axis, but on the distal side their axis switched (Fig. 1C). This indicates a change in orientation and alignment of the myofibers at the distal side.

**Figure 1.**
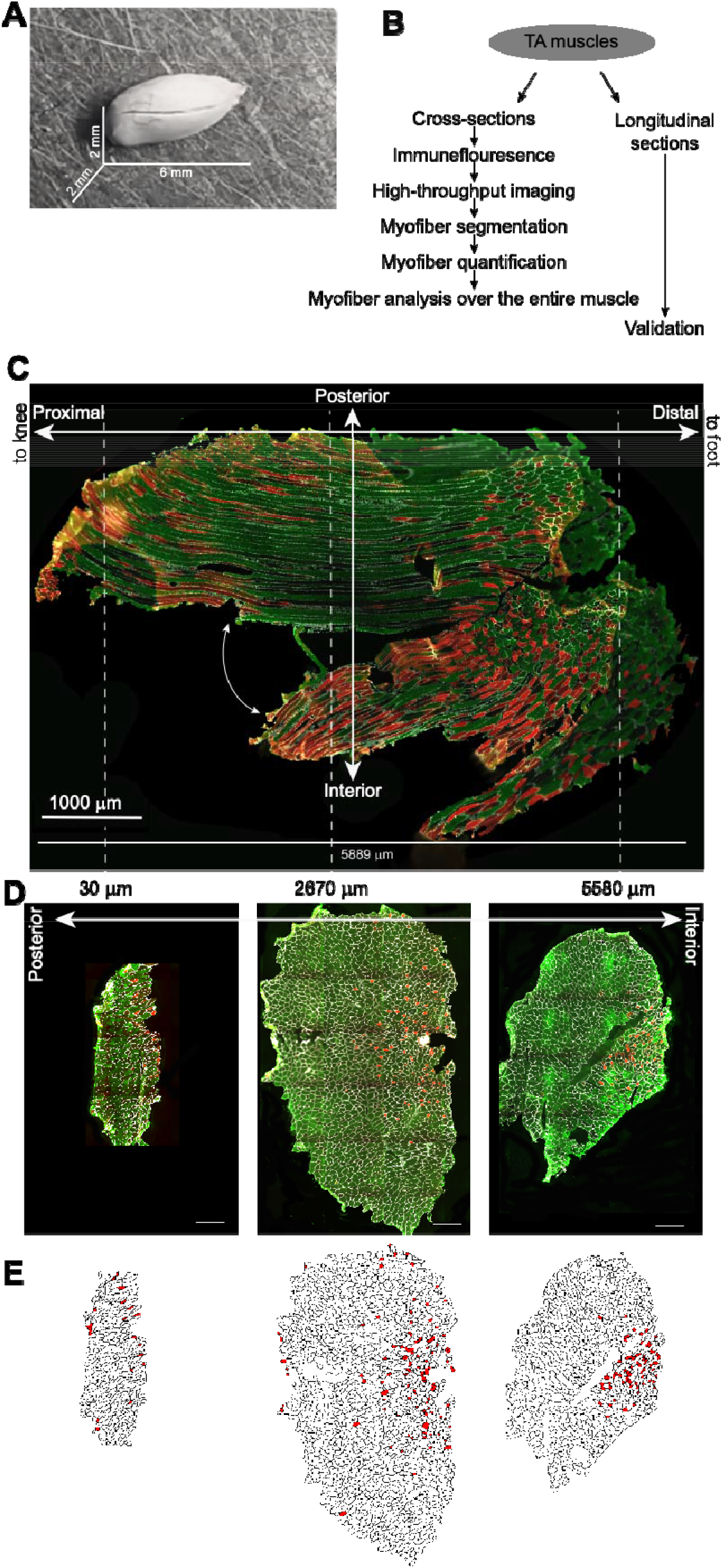
A summary of the experimental design and setup. **A)** An image of a TA muscle prior to sectioning. **B)** A methodological summary of myofiber data collection and analysis. **C)** An immunofluorescence image of a longitudinal section from a TA muscle. The section covers both distal and proximal ends at the longitudinal axis. The proximal end is to the knee and the distal end is to the foot. The interior and posterior ends are on the transverse axis, the interior side is towards the tibia. The entire muscle length is in μm. Scale bar is 1000 μm. Dashed white lines are estimates to proximal, median and distal regions. **D)** Immunofluorescence images of cross-sections from proximal and distal ends and from a median location. The distance of each cross-section from the proximal end is depicted above the image in μm. Cross-section orientation (Posterior – interior) is marked. Scale bar is 100 μm. **E)** Lamina segmentation of the cross-sections shown in D, with the MyHC-2A positive myofibers depicted in red. Immunofluorescence in C and D were carried out for Laminin (in white), MyHC-2A (in red), and MyHC-2B (in green).

To investigate changes in myofiber spatial organization along the longitudinal axis we aligned the cross-sections using the laminin-based segmentation and MyHC-2A positive fibers (Fig. 1E). The MyHC-2A myofibers were defined as positive when the mean fluorescence intensity (MFI) was greater than mean MFI + one standard deviation per section. Alignment of consecutive sections was performed using the scale-invariant feature transformation (SIFT), resulted in a good alignment (AVI. S1). But for an alignment of sections over the entire muscle, we had to compromise for a faster solution, in which the total data was reduced in size. As a guideline for data reduction, we allowed a gap of 150 μm between two consecutive sections, which smaller than the shortest myofiber length (160 μm). We kept this distance constant along the muscle to avoid a bias from over or under representative regions. With this criterion we ensured that all myofibers in the muscle tissue will be included in the analysis. In total, 129 (out of 150) cross-sections were used for consecutive alignment. The alignment outcome that were obtained with gaps between sections was also good and comparable to the alignment without gaps in the same region (AVI. S2, AVI S1, respectively). Since the alignment procedure only accounts for consecutive sections, consecutive alignment of pairs of sections (local) is hampered by the accumulation of errors. Therefore, we performed local alignments over parts of the muscle, and those gave good results for the proximal and median regions, but for the distal region alignments remained poor (Fig. S1). This further indicates that at the proximal and median regions myofibers have a similar parallel orientation but while those at the distal region have a multipennate orientation.

The change in myofiber alignment along the longitudinal axis suggests that myofiber properties might alter over the entire muscle. To investigate this, we considered only cryo-sections with good segmentation (>85% of the sections showed proper lamina segmentation). In total, 153,930 cross-section myofibers from 129 cross-sections were included in the analysis. From the segmented images, we extracted the myofiber cross-sectional area (CSA) and circularity. As the entire muscle was sectioned in a single session, the same cutting angle was kept along the entire muscle. Hence, a change in CSA or circularity could not reflect alternations in sectioning orientation but can indicate a biological phenomenon, such as curving, of muscle architecture. The change is each feature along the longitudinal axis is visualized with a smoothed regression line, which levels small variations between adjunct cross sections (Fig.2). This enables us assessing a global trend along the muscle, rather than the effect of an individual cross-section. The trend for the number of myofibers along the muscle longitudinal axis showed small numbers and higher number of myofibers in the middle of the muscle (Fig. 2A). This trend is expected from the macro shape of the TA muscle: smaller at both proximal and distal ends and larger in the centre of the muscle (Fig. 1A). The number of myofibers increased till 1/3^rd^ of the muscle’s length (position ~1800 μm) and overlapped with the position that marked a change in myofiber CSA (Fig. 2A-B). Myofiber median CSA was clearly smaller at the first ~1/3^rd^ of the muscle, and generally remains constant for the rest of the muscle. This proximal region, which can be discriminated from the rest of the muscle, was also observed using the median circularity: at the proximal region the median circularity was steadily increased (fibers become more circular), but slightly decreased in the other ~2/3^rd^ of the muscle (Fig. 2C). To assess the distribution of circularity scores, we calculated the variance of circularity and compared the median and variance curves along the entire muscle. Both curves showed opposite trends along the longitudinal axis (Fig. 2C). At the proximal end the circularity median was low, indicating an elongated orientation of the myofibers, and high variance, indicating heterogeneity. In contrast, at distal and median regions variance in circularity was low and median circularity was high (Fig. 2C). At the proximal side two sub-regions could be distinguished: 1)- low circularity and high variance, 2)- higher circularity and lower variance (Fig. 2C). In contrast, the changes in median and variance CSA along the muscle were similar (Fig. S2). We then overlaid the median CSA and median circularity curves in order to assess a spatial relation between the two features (Fig. 1D). With those two features, myofibers in the muscle sub-regions could be discriminated and described as following: at the proximal end (<1/9 of the muscle) myofibers CSA is small and circularity is low with high variance, subsequently circularity increases but myofiber CSA remains small (<1/9 of the muscle). The median part it the largest (~5/9^th^ of the muscle length), where myofibers have high CSA and circularity values but smaller variance in circularity (Fig. 2F). At the distal end, the relation between median CSA and mean circularity may slightly alter, this region also relates to the region with poor alignment (Fig. S1). Together, the analysis here revealed that geometrical features of myofibers changes along the longitudinal axis of the TA muscle.

**Figure 2.**
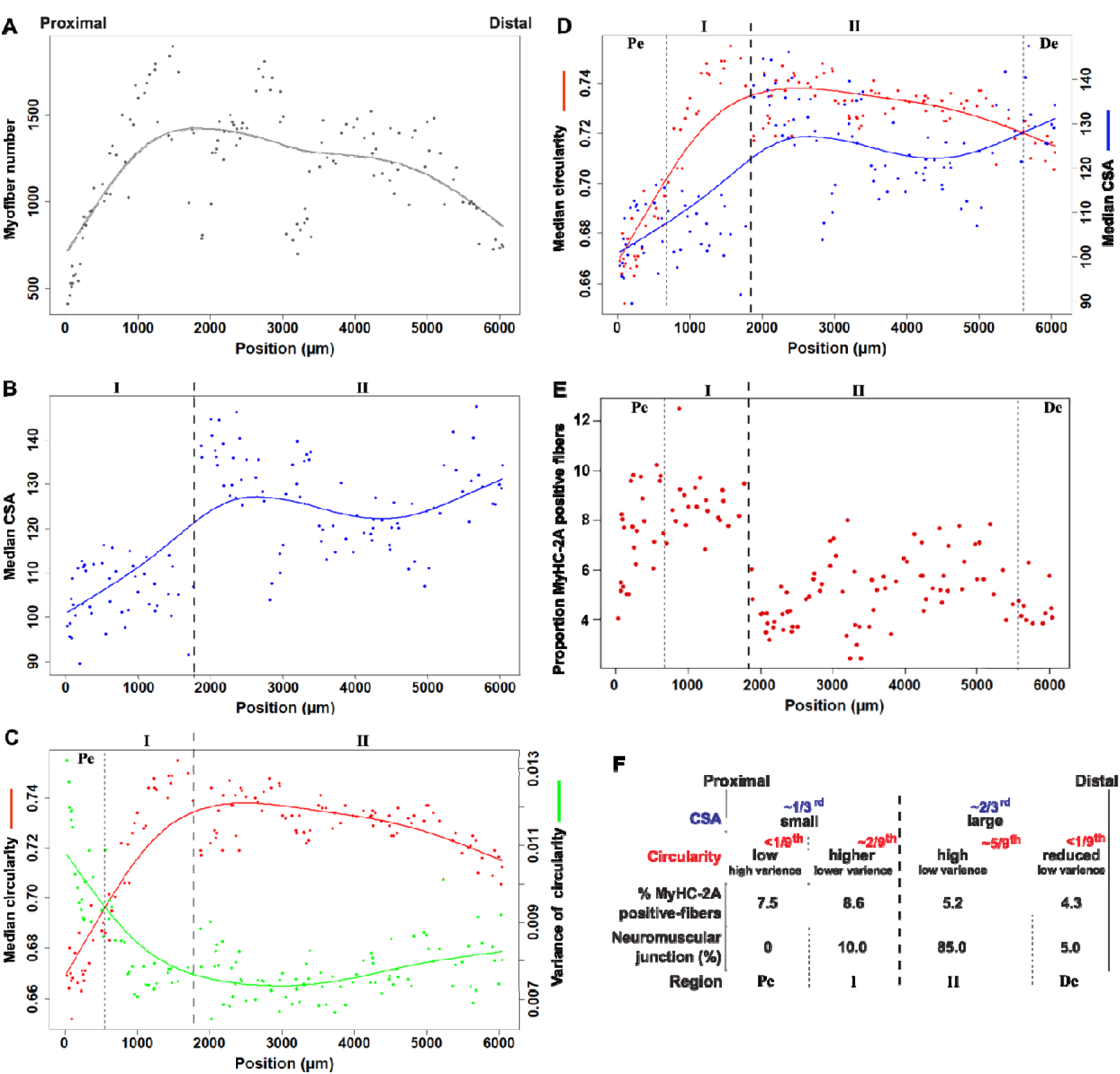
Spatial changes along the TA muscle. **A-E)** Scatter plots of measurements from each cross-section across the entire muscle. Every dot represents a cross-section. Cross-sections are positioned in a chronological order from the proximal to distal end (x-axis, in μm). A smoothed regression line is depicted. Vertical dashed line show muscle regions (I and II), the and dotted lines the proximal or distal ends (Pe and De, respectively). **A.** Number of myofibers normalized to the tissue area. **B.** Median of fiber CSA per section. **C.** Fiber circularity per section, mean (in red) and variance of the mean (in green). **D.** Mean circularity (in red) and median CSA (in blue). **E.** Proportion of MyHC-2A positive fibers. **F.** A summary of features per muscle region. Muscle regions that are based on the CSA trend are depicted in blue, and these made on circularity trends are in red.

We then investigated if these muscle regions, as defined by geometrical features of myofibers, can be distinguished by molecular features of muscle biology. We focus on the sarcomeric proteins, MyHC-2B and MyHC-2A, and the distribution of NMJs. The cross-sections were stained with antibody mix to MyHC isoforms, and after imaging the MFI were measured from each myofiber. Main myofiber-type subclasses were determined in the pooled dataset using density plots. The MyHC-2B density plot showed only a single peak, indicating a predominate one myofiber type (Fig. S3A), but two peaks were found in the MyHC-2A density plot (Fig. S3B). Based on the two peaks, we discriminated between the MyHC-2A positive myofibers and the MyHC-2A negative myofibers (Fig. S3B). We calculated the percentage of 2A-positive myofibers in every section (Fig. 2E) and in the muscle regions (Fig. S3B and Fig. 2F). Overall, the trend for the proportions of the 2A-positive myofibers is consistent with that of the median CSA: the proportions of 2A-positive myofibers were higher at the proximal region, compared with median and distal regions (Fig. 2E). The statistical significance of the differences between the percentages of the 2A-positive fibres in each region was significant (p-value < 2.2*10^−16^). The asymmetric distribution of MyHC-2A myofibers along the longitudinal axis suggests that contraction properties are not uniform along the muscle. Last, the NMJs were visualized in longitudinal sections with *α*-bungarotoxin, which specifically binds to NMJs (Tse *et al.*, 2014). A representative image is shown in (Fig. S4). Neuromuscular junctions were absent in the proximal-end and were hardly found in the distal end (Fig. 2F). The majority of NMJs were found in the medial region of the muscle.

## Discussion

Modelling of myofiber architecture in mouse TA muscles using non-invasive methods described a pennate architecture, where at the largest part of the muscle most fibers are parallel to the longitudinal axis and close to the tendon at the proximal and distal ends myofibers asymmetrically curve (Heemskerk *et al.*, 2005; Lovering *et al.*, 2013; Moo *et al.*, 2016). Here we performed ex-vivo analysis on the entire murine TA muscle, in both cross- and longitudinal-sections. In agreement with these studies, we also found asymmetric myofiber arrangement along the longitudinal axis, and between proximal and distal regions. At the proximal end myofibers were parallel to the longitudinal axis, but at the distal end myofiber orientation was altered, suggesting a bent structure. A bent structure of the TA muscle was indeed previously discussed (Moo *et al.*, 2016). The MyHC-2B positive myofibers seem to stretch over the entire muscle length, but the MyHC-2A positive myofibers are more abundant in the proximal part of the muscle. As both MyHC isoforms indicate different metabolic processes, their asymmetric distribution suggests differences in contraction properties along the muscle. Yet, we could not conclude whether the MyHC-2B myofibers at the distal end are the curved counterparts of the myofibers seen in the median part of the muscles.

In contrast to most studies that determine myofiber typing from a single region (often median), this study investigates the distribution of myofiber typing along the entire muscle. However, including all cross sections covering the entire muscle is highly laborious and requires an enormous computational power. Here we show that using the shortest myofiber length as a guideline for gaps between consecutive cross-sections enabled us to eliminate proportions of the cross-sections without losing spatial information. Since we show here that myofiber typing changes along the muscle, it potentially implicates that disease- and age-associated alternations in myofiber type composition could be more affected in specific muscle regions.

Using myofiber CSA and circularity we observed four regions along the TA muscle. Those muscle regions are also distinguished by the distribution of MyHC-2A positive myofibers and the NMJs. The largest muscle part is characterized by high circularity and high CSA, less MyHC-2A positive myofibers and higher density of NMJs. The shortest regions are the proximal and distal ends. Both differ in the orientation of myofibers (varied circularity), the percentage of MyHC-2A positive myofibers, and the proportions of NJMs. Based on circularity, myofibers at the proximal end are more heterogeneous compared to those at the distal end. An additional muscle (sub)region, next to the proximal end, could be defined based on geometrical features. This subregion could be a transition region between proximal and median regions. In this subregion NMJs were found, whereas at the proximal end no NMJs were found. Asymmetry of NMJ distribution along the muscle length has not been reported. However, it is recognized that NMJs higher NMJs concentration implicated higher contraction (Lomo, 2003).

In sum, despite the parallel organization of myofibers in the murine TA muscle, geometric, metabolic and molecular features seem to have an asymmetric distribution along the muscle. This is a pilot study and future studies with additional muscle markers are required to support the definition of muscle regions along the entire muscle. Moreover, more muscles are required to statistically assess the definition of our proposed myofiber regions.

## Additional Information

### Authors’ contribution

Wetlab experiments were performed by DB and CSO; imaging, image processing and image analysis were performed by DB and LMV; muscles were collected by MvP; project supervision was carried out by LMV, EvdA and VR; funding was received for VR; all authors wrote the MS.

## Supplementary material

**Figure S1.**
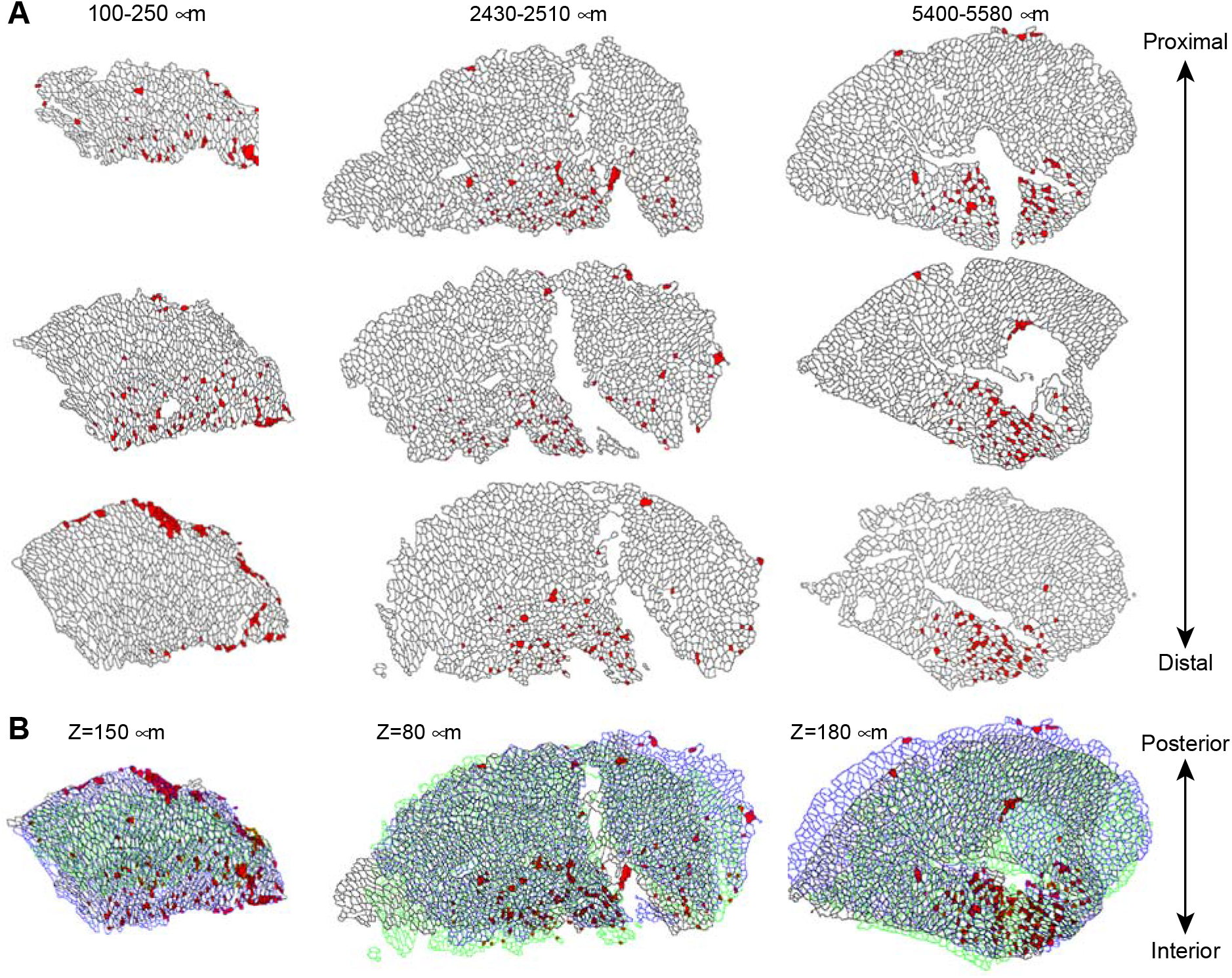
Alignment of three cross-sections in three muscle regions. A. Images of individual muscle cross-sections after fiber segmentation. The MyHC-2A-positive fibers are depicted in red. The range of the positions of the cross-sections is denoted above. B. Alignments of the three cross-sections. In the proximal and medial region, a good alignment was obtained, whereas alignment in the distal part was poor. The orientation of the cross-sections (proximal – distal, posterior – interior) is marked.

**Figure S2.**
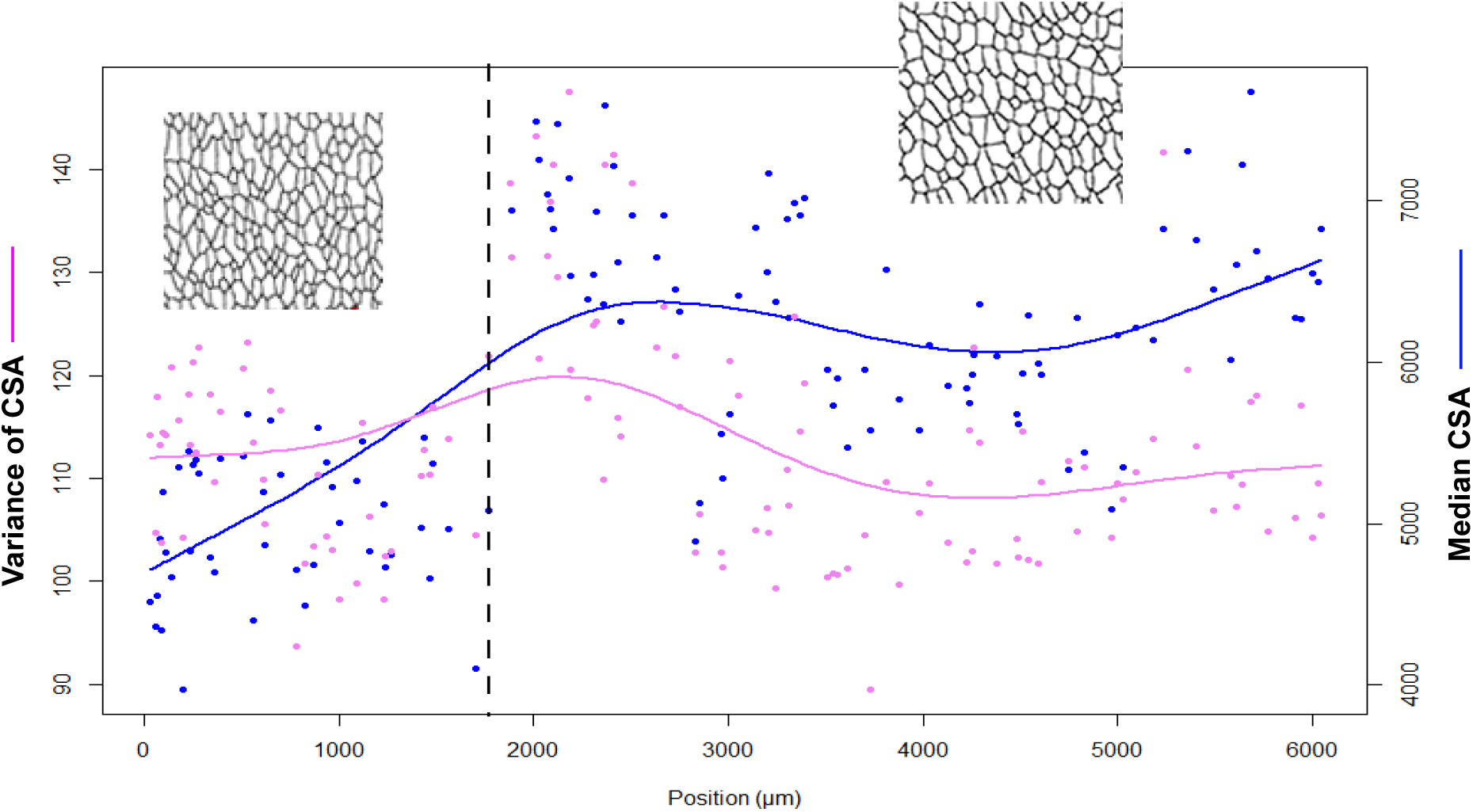
Scatter plot shows changes in median CSA and variance of the CSA along the entire muscle length. Each dot represents a cross-section, and the position of the cross-sections is marked on the X-axis (in μm). A representative lamina segmentation, for each muscle region is depicted above the scatter plot.

**Figure S3.**
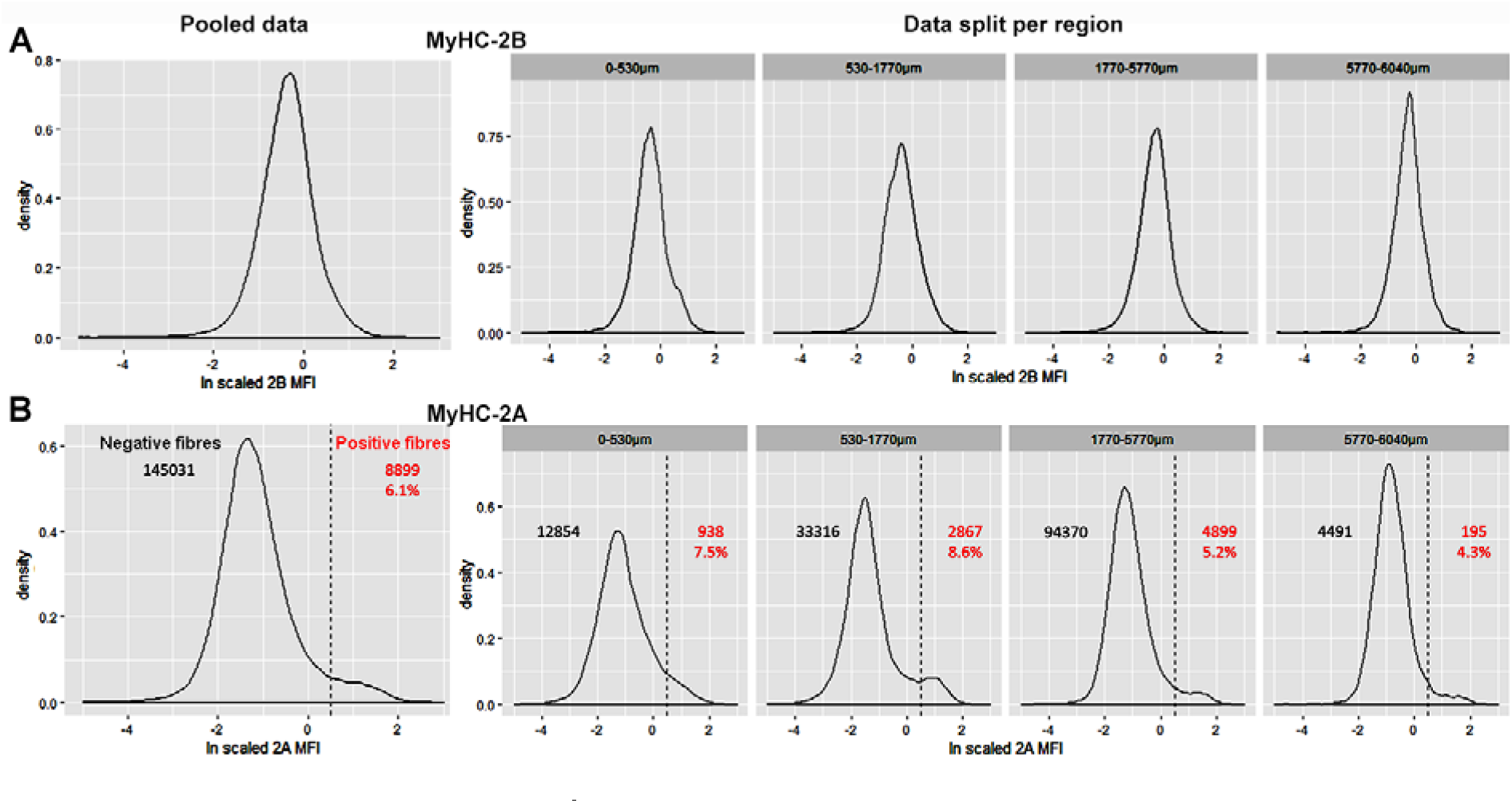
MFI density plots. The left panel shows MFI density in the pooled data, and the right panels show MFI density per muscle region, as defined in Figure 2. **A.** MyHC-2B. **B.** MyHC 2A; in each plot the number of 2A-negative or positive fibers are shown in black or red, respectively.

**Figure S4.**
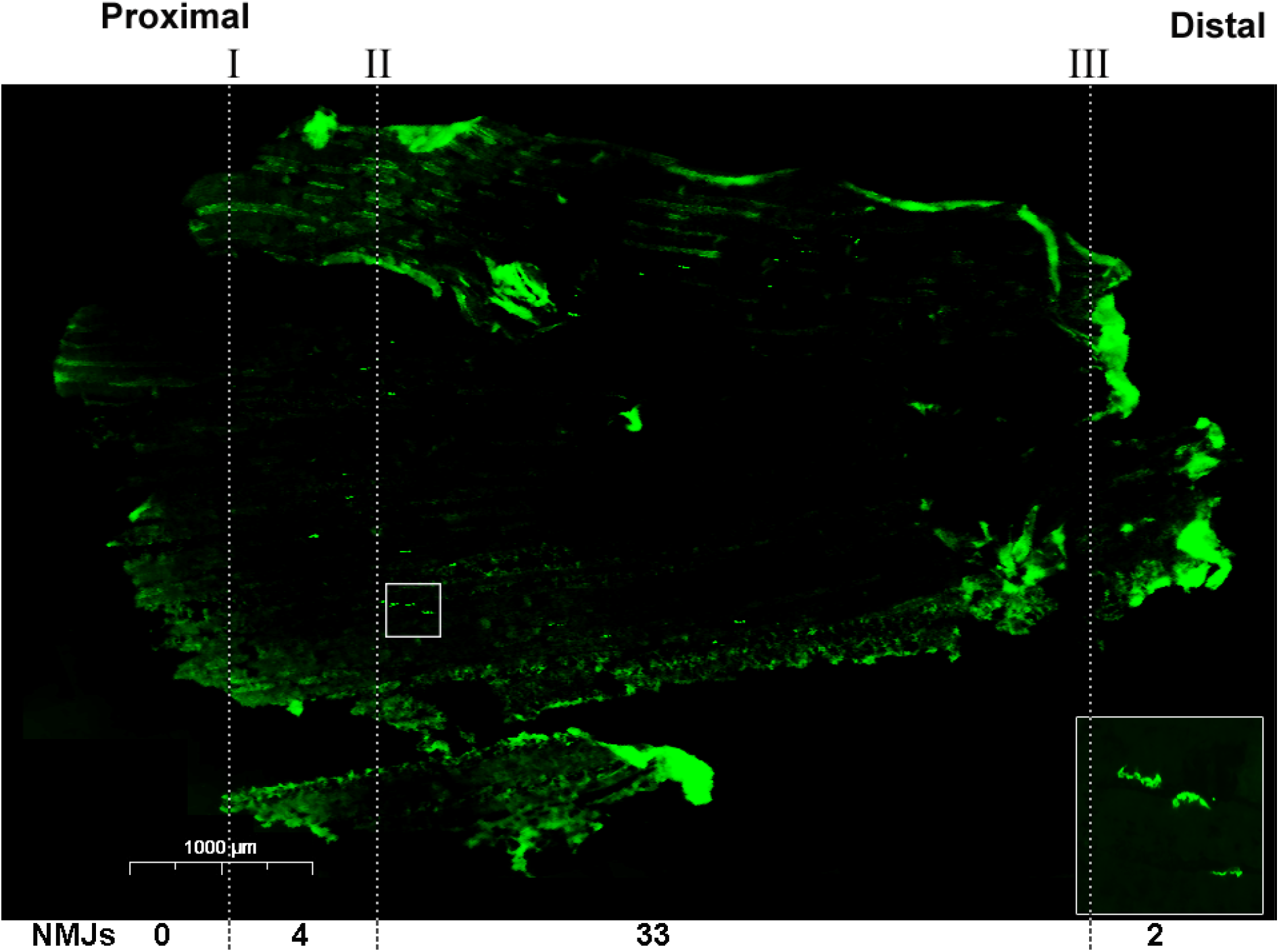
An immunofluorescence image of a longitudinal section stained with Alexa-488-conjugated α-bungarotoxin. In the white rectangle a magnification of staining of the NMJs on is depicted, from high magnification images the number of NMJs were counted. Scale bar is 1 mm. Dashed lines mark an estimate to muscle regions. The observed number of NMJs per muscle region are depicted under the photo.

**AVI S1.**
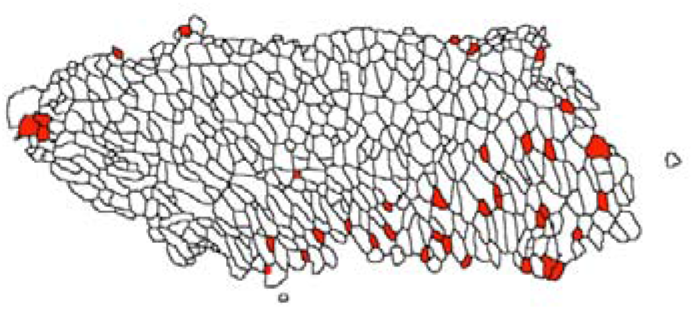
Animation of consecutive cross-sections alignment.

**AVI S2.**
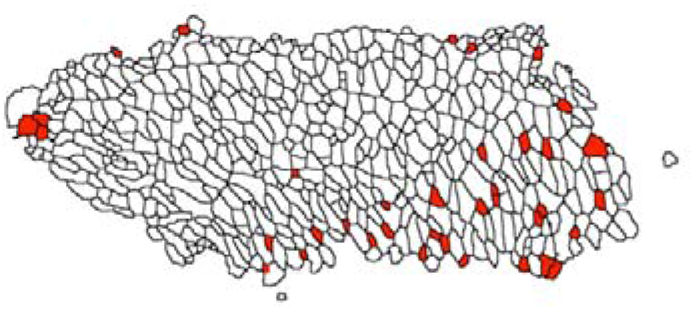
Animation of cross-sections alignment after 150 μm gapping.

